# Microsecond Time-Resolved Cryo-EM Based on Jet Vitrification

**DOI:** 10.1101/2025.11.21.689681

**Authors:** Michal Haubner, Harry M. Williams, Jakub Hruby, Monique S. Straub, Albert Guskov, Kirill Kovalev, Marcel Drabbels, Ulrich J. Lorenz

## Abstract

Understanding and ultimately predicting the function of proteins is one of the frontiers in structural biology. This will only be possible if it becomes feasible to routinely observe proteins on the fast timescales on which they perform their tasks. Recently, laser flash melting and revitrification experiments have improved the time resolution of cryo-electron microscopy (cryo-EM) to microseconds, rendering it fast enough to observe the domain motions of proteins that are frequently linked to function. However, observations have been limited to a time window of just a few hundred microseconds. Here, we introduce time-resolved cryo-EM experiments based on jet vitrification that combine microsecond resolution with an observation window of up to seconds. We use a short laser pulse to initiate protein dynamics, and as they unfold, vitrify the sample with a jet of a liquid cryogen to arrest the dynamics at that point in time. We demonstrate that our approach affords near-atomic spatial resolution and a time resolution of 21 µs. This allows us to observe the photoinduced dynamics of the light-driven sodium pump *Er*NaR on the microsecond to millisecond timescale. Our experiments significantly expand the ability of cryo-EM to observe protein dynamics across multiple timescales.

## Introduction

Understanding the dynamics of proteins is one of the frontiers in structural biology.^1,2^ Advances in instrumentation and computational tools have ushered in a golden age for protein structure determination,^3,4^ and the large number of structures that are now available have enabled machine learning approaches to even predict structures with good confidence.^3,5^ However, while a structure allows one to infer how a protein functions, one can frequently only do so imperfectly. Fully understanding and ultimately predicting protein function will only succeed if it becomes possible to routinely observe proteins as they perform their tasks, a goal that has so far remained out of reach.^1^ The development of new experimental methods is therefore urgently needed to advance our understanding of the dynamic nature of proteins.

X-ray crystallography has long been the workhorse of structural biology,^6^ and the introduction of high-brightness, pulsed X-ray sources has made it possible to observe protein dynamics with time-resolved protein crystallography.^7–11^ A number of experiments particularly stand out, in which the dynamics of light-activated proteins have been observed with ultrafast time resolution.^6,7,10^ However, the need for the sample to be crystalline limits the scope of the method, with the crystal often hindering some of the large-amplitude domain motions that are typically associated with the function of a protein.^1^

Time-resolved cryo-EM enables the observation of protein dynamics without the need for the particles to be confined to a crystal, but has traditionally offered only a limited time resolution.^12–18^ In these experiments, protein dynamics are typically initiated with a short pulse of light^19–21^ or by rapid mixing,^18,22^ after which the sample is vitrified to trap the particles in transient states. However, it has not been possible to plunge-freeze^20,21,23–26^ or jet vitrify^27,28^ samples with better than millisecond time-resolution, too slow to observe the microsecond domain motions that are frequently linked to function. We have recently introduced a new approach that makes the microsecond timescale accessible to cryo-EM.^29–31^ It involves flash melting a cryo-EM sample with a microsecond laser pulse to provide a brief time window, during which protein dynamics are initiated, before revitrification arrests the proteins in their transient configurations.^32–34^ However, observations with this technique have so far been limited to a time window of just a few hundred microseconds,^33^ leaving a gap to the millisecond timescale, which can already be accessed with traditional approaches. Ideally, one would like to cover all timescales in the same experiment. Here, we demonstrate time-resolved cryo-EM experiments based on jet vitrification that achieve this goal and combine microsecond time resolution with an observation window of up to seconds.

## Results and discussion

Figure 1a illustrates our experimental approach. A protein solution is applied to a specimen support, and dynamics are initiated with a short laser pulse, for example, by releasing a photocaged compound such as caged ATP, caged ions, or caged peptides^35–37^ Alternatively, conformational dynamics can be induced by creating a temperature jump^33^ or by directly exciting a photoactive protein, as we show below. As the dynamics unfold, the sample is vitrified with a precisely timed jet of a liquid cryogen to arrest the proteins in their transient configurations, which are subsequently imaged.

**Figure 1.**
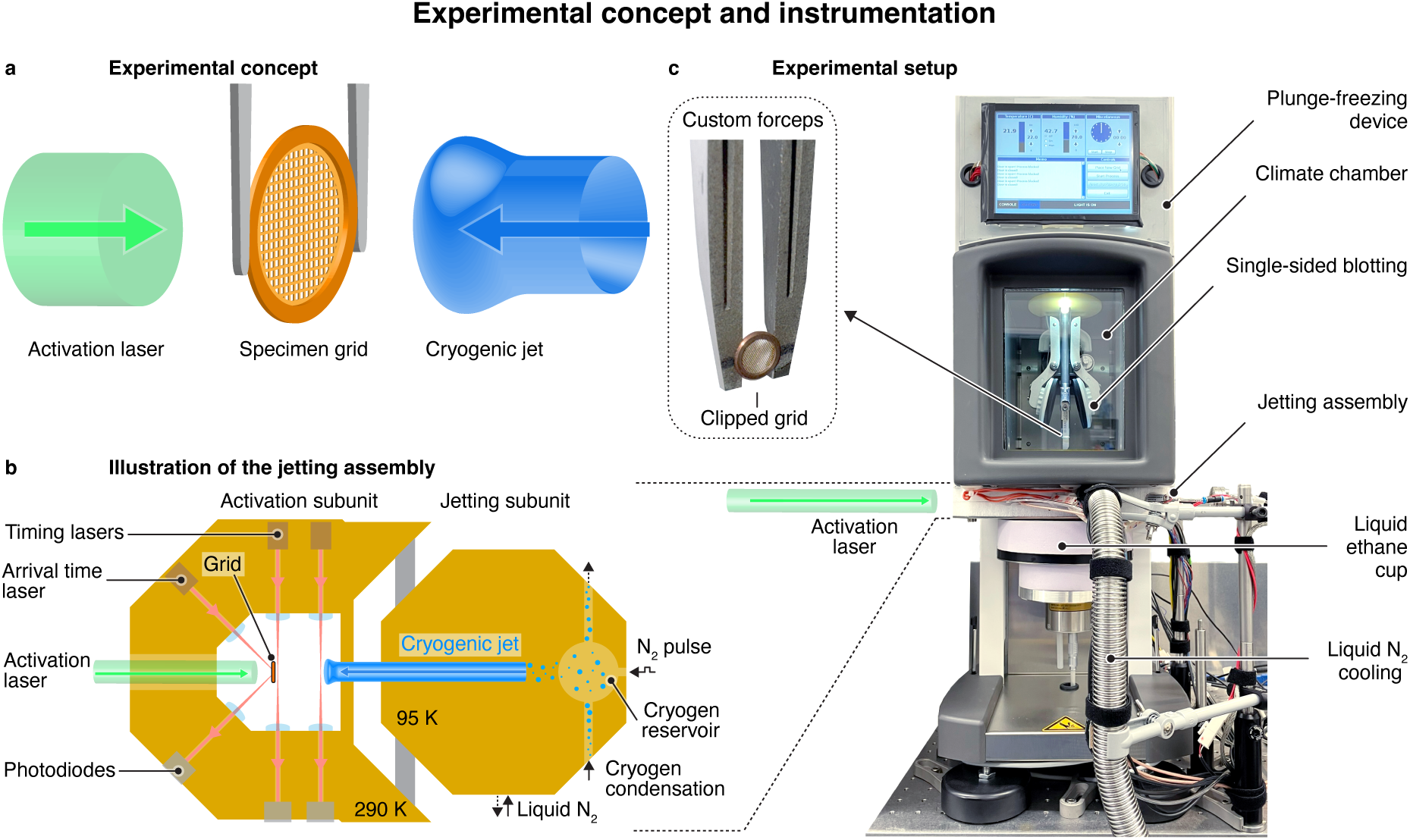
Experimental concept and instrumentation. **a** Experimental concept. Protein dynamics are initiated with a short laser pulse. As they unfold, the sample is vitrified with a time-delayed jet of a liquid cryogen, trapping the proteins in their transient configurations. **b** Illustration of the experimental implementation. A sample grid is placed in the activation subunit (left) of the jetting assembly, and protein dynamics are initiated with a laser pulse. The sample is then vitrified with a liquid ethane jet that is generated in the jetting subunit (right), where a small volume of the cryogen is condensed and subsequently expelled with a short pulse of compressed nitrogen. As the liquid ethane jet travels towards the specimen grid, its passage is detected at two points in the activation subunit with laser beams. This allows us to predict the arrival time of the jet on the fly and fire the activation laser such that the sample is vitrified at a desired time delay after dynamics have been initiated. The precise instant of vitrification is determined with a third laser beam that is continuously reflected off the back side of the grid and that is deflected by the arrival of the jet. **c** Experimental setup with the jetting assembly integrated into a commercial plunge freezing device (Thermo Fisher Vitrobot Mark IV). The sample is prepared in the climate chamber, using a clipped grid held in a custom forceps. After single-sided blotting, it is lowered into the jetting assembly, where the time-resolved jet vitrification experiment is performed. Finally, the specimen grid is lowered into a liquid ethane/propane filled cup underneath the jetting assembly.

Figure 1b illustrates our design of the time-resolved jet vitrification device (Supplementary Information 1). The specimen grid is positioned in the activation subunit on the left (held at a temperature of about 290 K), where it is illuminated with the activation laser pulse. The cryogenic jet is generated in the jetting subunit on the right, which is cooled to 95 K. A small volume of the cryogen (about 0.3 mL of ethane or an ethane/propane mixture) is condensed into a reservoir and subsequently expelled with a short pulse of compressed nitrogen. As the liquid jet travels towards the sample, which takes about 10.7 ms, we detect its passage at two points in the activation subunit with laser beams. When the jet passes these laser gates, the laser beams are deflected, which we detect with photodiodes. This allows us to predict the arrival time of the jet on the fly and fire the activation laser accordingly, so that the sample is vitrified at a desired time delay after dynamics have been initiated. We determine the precise moment at which the jet impinges on the sample with a third laser beam that is reflected off the back of the grid at an angle of 45°. The arrival of the jet deflects this laser beam, which we similarly detect with a photodiode.

As shown in the photograph of Fig. 1c, the jet vitrification device is integrated into a commercial plunge freezing apparatus (Thermo Fisher Vitrobot Mark IV) that we have modified for time-resolved experiments. The sample is prepared in the climate chamber of the plunge freezing device by applying a small volume of the protein solution onto a clipped specimen grid held in custom forceps (inset), followed by single-sided blotting. The sample is then lowered into the jetting assembly, which is mounted beneath the climate chamber, and the time-resolved jet vitrification experiment is performed. The specimen grid is then quickly lowered further into a liquid ethane/propane filled brass cup underneath the jetting assembly, from where it is transferred to an electron microscope for imaging.

Figure 2a shows high-speed video frames of a typical ethane jet impinging onto a sample (Supplementary Information 2). The jet, traveling at a speed of 2.8 m/s, has a diameter of approximately 3 mm and features a slightly curved front with a radius of curvature of about 1.9 mm at its center. Upon contact with the jet, the sample cools rapidly. Figure 2b shows a typical temperature evolution (red), as measured with a resistance temperature detector placed at the sample location (Supplementary Information 3, 1:1 ethane-propane jet with a speed of 2.65 m/s). An exponential fit (solid black line) yields a 1/e cooling time of 15.7 μs, which provides an estimate of how rapidly we can induce vitrification and trap proteins in their transient configurations. Figure 2c confirms that we can obtain near-atomic resolution reconstructions from jet vitrified samples, with a reconstruction of apoferritin yielding a resolution of 1.3 Å (Supplementary Information 5–6).

**Figure 2.**
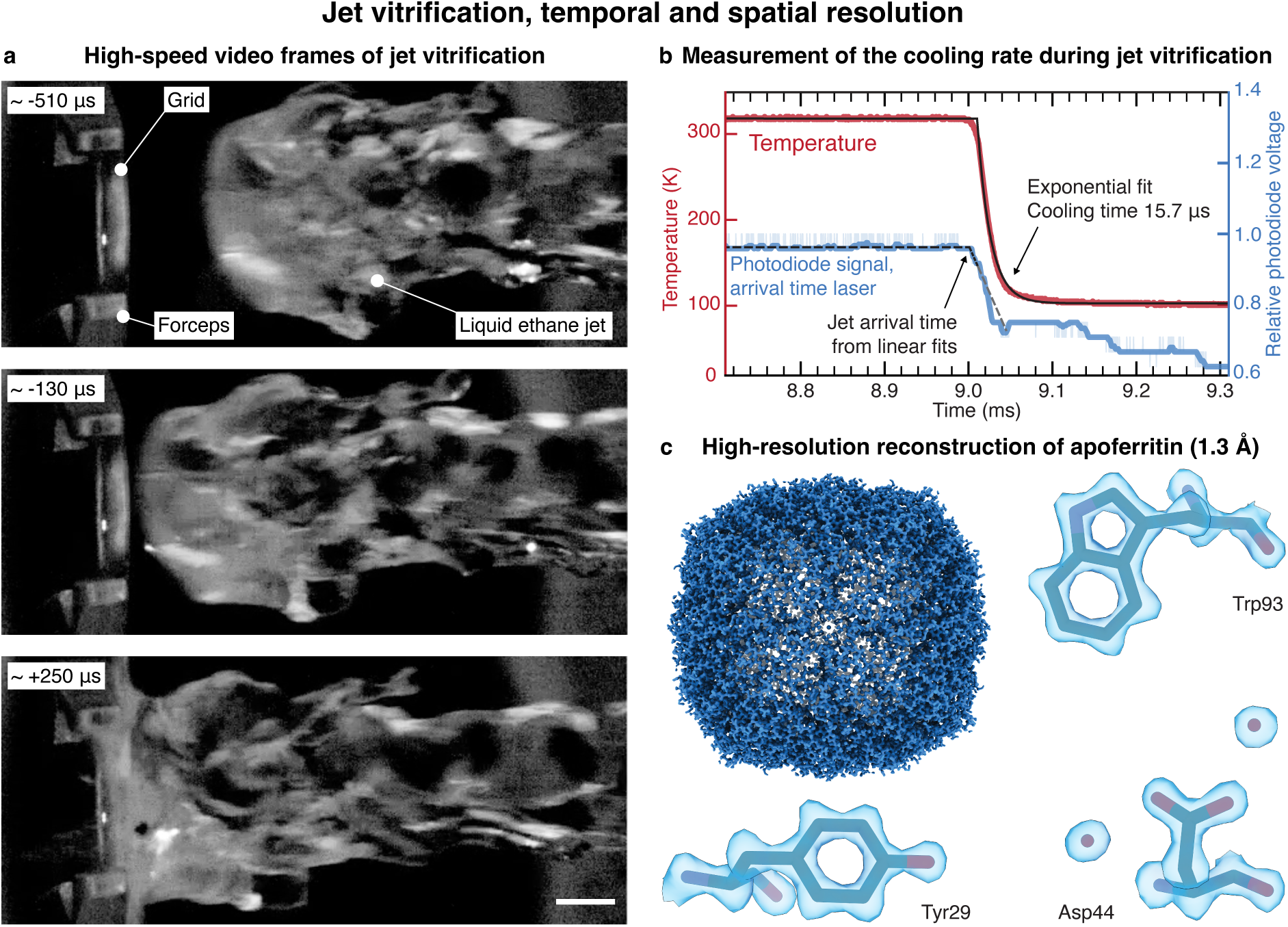
High-speed video frames of the jet vitrification process as well as characterization of the temporal and spatial resolution. **a** High-speed video frames of a liquid ethane jet, traveling at a speed of about 2.8 m/s, as it impinges on the sample. The time stamp of each frame is indicated with respect to the arrival time of the jet. Scale bar, 1 mm. **b** Typical temperature evolution (red) during the impact of the jet (1:1 ethane-propane, traveling at 2.65 m/s), as measured with a resistance temperature detector. An exponential fit (solid black line) yields a 1/e cooling time of 15.7 μs. The impact of the jet is detected with the arrival time laser, which is reflected off the back of temperature detector. As the jet impinges on the detector, the laser beam is deflected, which is detected with a photodiode (light blue, the dark blue curve represents the smoothed signal). We determine the timing of this event from the intersection of two linear fits to the signal, which are shown with dashed, black lines (the dashed, grey line provides a guide to the eye). The time is indicated with respect to the trigger sent to the pulsed valve. **c** A reconstruction of apoferritin from a jet vitrified sample yields a resolution of 1.3 Å. A model is placed into the density with rigid-body fitting (PDB: 8RQB).^38^

We precisely time the cryogenic jet with respect to the excitation laser pulse, which allows us to perform experiments with microsecond time resolution. If we synchronize the activation laser to the pulse of compressed nitrogen that expels that cryogenic jet, we can realize arbitrarily long time delays, but only with an accuracy of 0.78 ms (as measured by the standard deviation, Supplementary Information 4). This is due to the jitter in the opening of the pulsed valve as well as variations in the jet speed. We can improve the accuracy to 0.39 ms by detecting the passage of the jet with the first laser gate and triggering the activation laser accordingly. This allows us to realize time delays of up to ~3 ms, the time it takes for the jet to travel from the first laser gate to the sample. For even shorter time delays of up to ~300 µs, we detect the jet with both laser gates to predict its arrival time on the fly, which yields an accuracy of 38 μs. Note that this experimental design allows us to time the jet more precisely for shorter time delays. This reduces the number of experiments that have to be rejected because the relative timing error is too large, *i.e.* the relative difference between actual and desired time delay. We determine the actual arrival time of the jet with the laser beam reflected off the back of the specimen grid (Supplementary Information 4). As shown in Fig. 2b, the arrival of the jet deflects the laser beam, which we detect with a photodiode (light blue, with the smoothed signal shown in dark blue and linear fits indicated with dashed, black lines). This allows us to determine the arrival time with an accuracy of 9 µs, independently of the chosen time delay. If we also take the average 1/e cooling time of the sample of 19 μs into account, we can estimate that our jet vitrification experiment has a time resolution of 21 µs (Supplementary Information 3). Note that because of the curvature of the jet front (Fig. 2a), sample areas further away from the center of the grid likely vitrify later. In the time-resolved experiments presented below, we have therefore only collected micrographs from the center of the grid (Supplementary Information 6–7).

We demonstrate the capabilities of our method by studying the dynamics of the rhodopsin *Er*NaR (*Erythrobacter sp*. HL-111), which upon light activation, pumps sodium ions from the cytoplasm across the bacterial membrane to the ouside.^39^ *Er*NaR belongs to a recently identified subgroup of the NDQ rhodopsins, which feature an additional glutamic acid residue close to the retinal Schiff base, allowing *Er*NaR to efficiently pump sodium ions even at acidic pH.^39^ Uncovering the structural dynamics of *Er*NaR therefore promises to shed light on the sodium pumping mechanism of this subgroup. The dark state structure of the pentameric *Er*NaR is shown in Fig. 3a (Supplementary Information 5, 7, as derived from a 2.1 Å resolution map), and the proposed ion transport pathway is illustrated.^39^ The photocycle with its relevant timescales is shown in Fig. 3b.^39^ Rhodopsins and other retinal-containing proteins have been popular targets for time-resolved X-ray crystallography studies.^40–44^ Although we cannot match the ultrafast time resolution of these experiments and therefore miss some of the earliest events that follow the photoisomerization of the retinal Schiff base, our jet vitrification experiments can be performed with a benchtop instrument, using orders of magnitude less sample than required for time-resolved serial crystallography.^11^ Moreover, we are able to study *Er*NaR in its naturally occurring pentameric form, which so far has resisted crystallization.^39^

**Figure 3.**
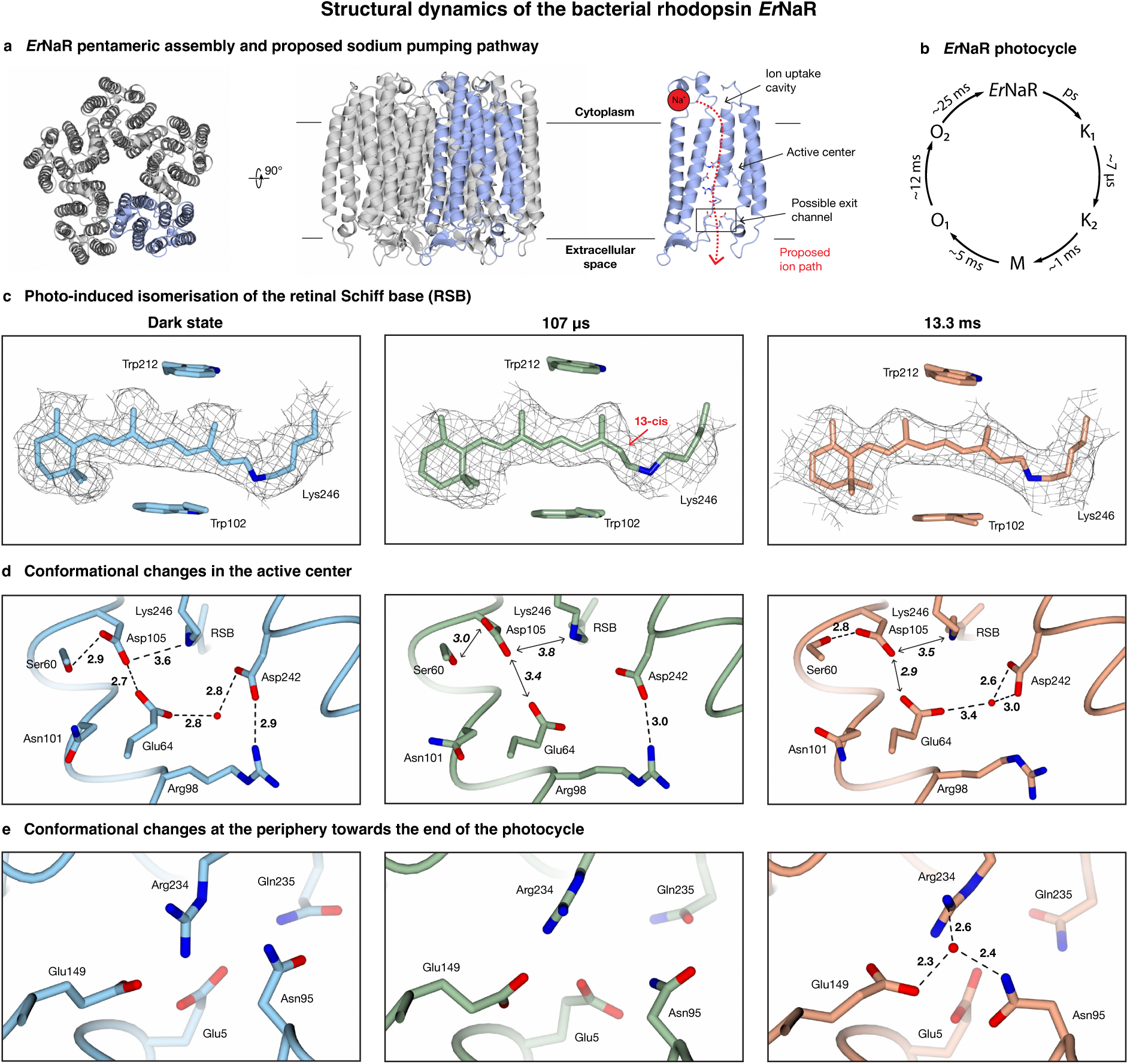
Structural dynamics of the bacterial rhodopsin *Er*NaR observed with microsecond time-resolved cryo-EM based on jet vitrification. **a** Dark state of the pentameric *Er*NaR, with a single monomer highlighted in blue, and proposed pathway along which the sodium ion is transported upon light activation (red, shown in a cutaway view of a monomer). **b** Illustration of the photocycle with the relevant timescales indicated.^39^ Note that these timescales were determined for a 1000 mM sodium ion concentration, which is higher than the concentrations used in our experiment (Supplementary Information 5). However, this is not expected to significantly change the timescales of the states we probe. **c,d,e** Structural changes observed at 107 µs and 13.3 ms compared to the dark state. Light absorption leads to the isomerization of the retinal Schiff base (RSB), which returns to the all-*trans* on long timescales (**c**). In the active center close to the retinal Schiff base, changes in the hydrogen bonding pattern suggest the uptake and transport of sodium ions (**d**). At long times, a putative sodium binding site forms near the exit of the ion transport channel, from which the ion is likely released to the extracellular space (**e**), in analogy to KR2.^40^ Hydrogen bonds are indicated with dashed lines, with the bond lengths given in Å. Double headed arrows indicate disrupted interactions.

Samples of *Er*NaR are activated with a nanosecond laser pulse of 532 nm wavelength (6 ns FWHM pulse duration, 6.4 mJ/cm^2^ fluence) and jet vitrified at 107 µs and 13.3 ms, to probe the K_2_ and O_2_ states, respectively (Supplementary Information 5, 7). We analyze the structural dynamics of *Er*NaR at the level of a monomer (C_5_ symmetry expansion), neglecting potential cooperativity between the subunits because of the small amplitude of the motions involved. We remove monomers that are not undergoing the photocycle with 3D classification, which then allows us to obtain a 2.8 Å and 3.3 Å reconstruction for the K_2_ and O_2_ state, respectively, and build molecular models (Supplementary Information 7). Figure 3c–e illustrates key features of the structural dynamics occurring in the retinal Schiff base, in the active center close to it, as well as in a site near the exit of the ion transport channel (highlighted in Fig. 3b), where sodium has been proposed to transiently bind in the O_2_ state of monomeric KR2,^40^ a well-studied sodium pump.

At 107 µs after photoactivation, we observe the K_2_ state. As shown in Fig. 3c, the chromophore has been switched from the all-*trans* configuration present in the dark state to the 13-*cis* configuration,^45,46^ a movement that is accompanied by small rotations of the aromatic sidechains in the retinal binding pocket. Moreover, in the adjacent active center, a disruption of the tight hydrogen bond network occurs that characterizes the dark state (Fig. 3d). This network, which includes Lys246 to which the retinal cofactor is covalently bound, is believed to prevent transport through the channel. Photoisomerization of the chromophore is thought to partially disrupt this hydrogen bonding network allowing a sodium ion to be translocated through the cavity of the active center.^39,47^ In particular, the densities for the side chains of Glu64 and Asp105 suggest increased flexibility. In the dark state, these sidechains are strongly connected through a short hydrogen bond, which has been suggested to provide the structural basis for allowing *Er*NaR to efficiently pump sodium even at acidic pH. Here, this hydrogen bond appears to have been disrupted, along with the hydrogen bond between the retinal Schiff base and Asp105. Moreover, the density for the water molecule that is coordinated by Glu64 and Asp242 in the dark state is absent. We propose that these conformational changes are related to priming of the active center for the translocation of sodium.

At 13.3 ms, we observe *Er*NaR in the O_2_ state. The O states are associated with the sodium ion translocation, but differ in the configuration of the retinal Schiff base.^48,49^ Transient absorption spectroscopy suggests that the transition from the O_1_ to the O_2_ state is accompanied by an isomerization of the chromophore from the 13-*cis* back to the all-*trans* configuration.^39^ Indeed, Fig. 3d shows that the retinal Schiff base adopts the all-*trans* conformation, with the aromatic sidechains surrounding the retinal returned to their dark state positions. In the active center, the hydrogen bonding network has partially reformed (Fig. 3e). The sidechains of Asp105 and Ser60 now show a stable interaction, and Glu64 and Asp242 coordinate a water molecule. These conformational changes potentially indicate transient sodium binding near the retinal Schiff base, as observed for the pentameric sodium pump KR2.^50^ Finally, we also observe conformational dynamics at a site near the exit of the ion transport channel (Fig. 3e), which for monomeric KR2, has been proposed to be a transient sodium binding site in the O_2_ state.^40^ The sidechain of Asn95 rotates into the binding site, which could feasibly facilitate transient sodium ion binding. We observe a spherical density at this position, which we cannot unambiguously identify as a sodium ion due to the limited resolution of the map and which we have therefore chosen to model as a water molecule. It appears likely that our map contains multiple transient sodium binding configurations, which limits its resolution, allowing only speculative interpretation at this stage.

## Conclusion

In conclusion, we have demonstrated microsecond time-resolved cryo-EM experiments based on jet vitrification. Our method significantly expands the ability of cryo-EM to observe protein dynamics and presents an alternative to flash melting and revitrification experiments for achieving microsecond time resolution.^31^ With further technical improvements, it should be possible to time the cryogenic jet even more accurately and ultimately match the few-microsecond time resolution afforded by flash melting experiments. Jet vitrification offers the advantage that it does not suffer from a limited temporal observation window, allowing us to observe protein dynamics continuously from a timescale of microseconds to seconds. Moreover, the sample temperature can be controlled easily, whereas in flash melting experiment, the sample follows a complex temperature evolution.^33,51^ For now, our experiments are limited to processes that can be initiated with a laser pulse, for example by releasing a photocaged compound^35–37^, inducing a temperature jump, or activating a light-sensitive protein, as we have done here. To broaden the range of dynamics that can be studied, it will therefore be crucial to develop approaches that enable the mixing of two reactants on the grid with microsecond time resolution.

Finally, we expect that our experimental approach should find applications beyond studying single-particle dynamics. For example, we have recently shown that if we replace laser excitation with mechanical excitation of the sample by an ultrasonic transducer during jet vitrification, we are able to overcome preferred orientation, even for such a notoriously challenging protein as hemagglutinin.^52^ Jet vitrification has also been shown to enable the vitrification of thicker objects, including cells,^28^ which suggests opportunities for performing microsecond time-resolved cryo-electron tomography experiments with our experimental design. It should also be possible to employ our approach to prepare freeze-quenched samples with microsecond time resolution for other techniques than cryo-EM, such as EPR spectroscopy, X-ray absorption, optical spectroscopy, or FRET experiments.

## Author contributions

Conceptualization: MH, UJL, MD

Methodology: MH, JH, UJL, HMW, AG, KK

Investigation: MH, JH, MSS, HMW

Visualization: MH, JH, MSS, HMW, KK

Funding acquisition: UJL, AG, KK

Project administration: UJL, AG, KK

Supervision: UJL, MD

Writing – original draft: MH, JH, MSS, HMW

Writing – review & editing: MH, JH, MSS, UJL, MD, HMW, AG, KK

## Data and materials availability

Coordinates and maps included in this paper have been deposited in the Worldwide Protein Data Bank (wwPDB) and the Electron Microscopy Data Bank (EMDB) with the following accession codes: apoferritin EMD-55761; *Er*NaR pentamer [dark state] EMD-55765 PDB 9TBD; *Er*NaR monomer [active state – 107 µs] EMD-55766 PDB 9TBE; *Er*NaR monomer [active state – 13.3 ms] EMD-55769 PDB 9TBF. The micrographs were deposited on the Electron Microscopy Public Image Archive (EMPIAR).

## Acknowledgments

The authors would like to thank the EPFL ISIC electrical and mechanical workshops for their help in realizing this project, in particular S. Dutoit, B.C. Le Geyt, and G. Pasche. The authors would also like to thank the EPFL Protein Production and Structure Core Facility for their help producing the apoferritin used in this project, and the Kikkawa Lab for making the apoferritin expression plasmid available.^53^ Cryo-EM data collection was performed at the Dubochet Center for Imaging Lausanne (a joint initiative from EPFL, UNIGE, UNIL, UNIBE) with the assistance of A. Myasnikov, B. Beckert, S. Nazarov, I. Mohammed, and E. Uchikawa.

## References

1. Henzler-Wildman, K. & Kern, D. Dynamic personalities of proteins. Nature 450, 964–972 (2007).

2. Ourmazd, A., Moffat, K. & Lattman, E. E. Structural biology is solved — now what? Nat. Methods 19, 24–26 (2022).

3. Jumper, J. & Hassabis, D. Protein structure predictions to atomic accuracy with AlphaFold. Nat. Methods 19, 11–12 (2022).

4. Kühlbrandt, W. Biochemistry. The resolution revolution. Science 343, 1443–1444 (2014).

5. Baek, M. et al. Accurate prediction of protein structures and interactions using a three-track neural network. Science 373, 871–876 (2021).

6. Shi, Y. A glimpse of structural biology through X-ray crystallography. Cell 159, 995–1014 (2014).

7. Schotte, F. et al. Watching a protein as it functions with 150-ps time-resolved x-ray crystallography. Science 300, 1944–1947 (2003).

8. Christou, N.-E. et al. Time-resolved crystallography captures light-driven DNA repair. Science 382, 1015–1020 (2023).

9. Banari, A. et al. Advancing time-resolved structural biology: latest strategies in cryo-EM and X-ray crystallography. Nat. Methods 22, 1420–1435 (2025).

10. Pande, K. et al. Femtosecond structural dynamics drives the trans/cis isomerization in photoactive yellow protein. Science 10.1126/science.aad5081 (2016) doi:10.1126/science.aad5081.

11. Brändén, G. & Neutze, R. Advances and challenges in time-resolved macromolecular crystallography. Science 373, eaba0954 (2021).

12. Mäeots, M.-E. & Enchev, R. I. Structural dynamics: review of time-resolved cryo-EM. Acta Crystallogr. Sect. Struct. Biol. 78, 927–935 (2022).

13. Frank, J. Time-resolved cryo-electron microscopy: Recent progress. J. Struct. Biol. 200, 303–306 (2017).

14. Klebl, D. P., Kay, R. W., Sobott, F., Kapur, N. & Muench, S. P. Towards sub-millisecond cryo-EM grid preparation. Faraday Discuss. 240, 33–43 (2022).

15. Klebl, D. P. et al. Need for Speed: Examining Protein Behavior during CryoEM Grid Preparation at Different Timescales. Structure 28, 1238–1248.e4 (2020).

16. Huber, S. T. et al. Nanofluidic chips for cryo-EM structure determination from picoliter sample volumes. eLife 11, e72629.

17. Torino, S., Dhurandhar, M., Stroobants, A., Claessens, R. & Efremov, R. G. Time-resolved cryo-EM using a combination of droplet microfluidics with on-demand jetting. Nat. Methods 20, 1400–1408 (2023).

18. Klebl, D. P., White, H. D., Sobott, F. & Muench, S. P. On-grid and in-flow mixing for time-resolved cryo-EM. Acta Crystallogr. Sect. Struct. Biol. 77, 1233–1240 (2021).

19. Shaikh, T. R., Barnard, D., Meng, X. & Wagenknecht, T. Implementation of a flash-photolysis system for time-resolved cryo-electron microscopy. J. Struct. Biol. 165, 184–189 (2009).

20. Bhattacharjee, B., Rahman, M. M., Hibbs, R. E. & Stowell, M. H. B. A simple flash and freeze system for cryogenic time-resolved electron microscopy. Front. Mol. Biosci. 10, 1129225 (2023).

21. Yoder, N. et al. Light-coupled cryo-plunger for time-resolved cryo-EM. J. Struct. Biol. 212, 107624 (2020).

22. Alexandrescu, L., Lessin, W. & Lander, G. C. Mix-it-up: Accessible time-resolved cryo-EM on the millisecond timescale. 2025.07.22.666177 Preprint at 10.1101/2025.07.22.666177 (2025).

23. Shaikh, T. R., Barnard, D., Meng, X. & Wagenknecht, T. Implementation of a flash-photolysis system for time-resolved cryo-electron microscopy. J. Struct. Biol. 165, 184–189 (2009).

24. Montaño Romero, A., Bonin, C. & Twomey, E. C. C-SPAM: an open-source time-resolved specimen vitrification device with light-activated molecules. IUCrJ 11, 16–22 (2024).

25. Levitz, T. S. et al. Approaches to Using the Chameleon: Robust, Automated, Fast-Plunge cryoEM Specimen Preparation. Front. Mol. Biosci. 9, (2022).

26. Dandey, V. P. et al. Time-resolved cryoEM using Spotiton. Nat. Methods 17, 897–900 (2020).

27. Henderikx, R. J. M. et al. VitroJet: new features and case studies. Acta Crystallogr. Sect. Struct. Biol. 80, 232–246 (2024).

28. Gemin, O. et al. EasyGrid: A versatile platform for automated cryo-EM sample preparation and quality control. 2024.01.18.576170 Preprint at 10.1101/2024.01.18.576170 (2024).

29. Bongiovanni, G., Harder, O. F., Drabbels, M. & Lorenz, U. J. Microsecond melting and revitrification of cryo samples with a correlative light-electron microscopy approach. Front. Mol. Biosci. 9, 1044509 (2022).

30. Klebl, D. P., Aspinall, L. & Muench, S. P. Time resolved applications for Cryo-EM; approaches, challenges and future directions. Curr. Opin. Struct. Biol. 83, 102696 (2023).

31. Lorenz, U. J. Microsecond time-resolved cryo-electron microscopy. Curr. Opin. Struct. Biol. 87, 102840 (2024).

32. Harder, O. F., Barrass, S. V., Drabbels, M. & Lorenz, U. J. Fast viral dynamics revealed by microsecond time-resolved cryo-EM. Nat. Commun. 14, 5649 (2023).

33. Curtis, W. A. et al. Ultrathin Liquid Cells for Microsecond Time-Resolved Cryo-EM. 2025.05.05.652279 Preprint at 10.1101/2025.05.05.652279 (2025).

34. Barrass, S. V. et al. Cryo-EM Sample Preparation with Soft-Landing and Laser Flash Melting. 2025.06.05.657968 Preprint at 10.1101/2025.06.05.657968 (2025).

35. Gutman, M. Gutman: Methods of Biochemical Analysis Ch. 1. (John Wiley & Songs, Inc., New York, 1984).

36. Shigeri, Y., Tatsu, Y. & Yumoto, N. Synthesis and application of caged peptides and proteins. Pharmacol. Ther. 91, 85–92 (2001).

37. Ellis-Davies, G. C. R. Caged compounds: photorelease technology for control of cellular chemistry and physiology. Nat. Methods 4, 619–628 (2007).

38. Küçükoğlu, B. et al. Low-dose cryo-electron ptychography of proteins at sub-nanometer resolution. Nat. Commun. 15, 8062 (2024).

39. Podoliak, E. et al. A subgroup of light-driven sodium pumps with an additional Schiff base counterion. Nat. Commun. 15, 3119 (2024).

40. Skopintsev, P. et al. Femtosecond-to-millisecond structural changes in a light-driven sodium pump. Nature 583, 314–318 (2020).

41. Wranik, M. et al. A multi-reservoir extruder for time-resolved serial protein crystallography and compound screening at X-ray free-electron lasers. Nat. Commun. 14, 7956 (2023).

42. Panneels, V. et al. Time-resolved structural studies with serial crystallography: A new light on retinal proteins. Struct. Dyn. 2, 041718 (2015).

43. Grünbein, M. L. et al. Crystallographic Studies of Rhodopsins: Structure and Dynamics. in Rhodopsin: Methods and Protocols (ed. Gordeliy, V.) 147–168 (Springer US, New York, NY, 2022). doi:10.1007/978-1-0716-2329-9_7.

44. Caramello, N. & Royant, A. From femtoseconds to minutes: time-resolved macromolecular crystallography at XFELs and synchrotrons. Acta Crystallogr. Sect. Struct. Biol. 80, 60–79 (2024).

45. Morizumi, T. et al. Structural insights into light-gating of potassium-selective channelrhodopsin. Nat. Commun. 16, 1283 (2025).

46. Lamm, G. H. U. et al. CryoRhodopsins: a comprehensive characterization of a group of microbial rhodopsins from cold environments. 2024.01.15.575777 Preprint at 10.1101/2024.01.15.575777 (2024).

47. Kovalev, K. et al. Structure and mechanisms of sodium-pumping KR2 rhodopsin. Sci. Adv. 5, eaav2671 (2019).

48. Inoue, K. et al. A light-driven sodium ion pump in marine bacteria. Nat. Commun. 4, 1678 (2013).

49. Gushchin, I. et al. Crystal structure of a light-driven sodium pump. Nat. Struct. Mol. Biol. 22, 390–395 (2015).

50. Kovalev, K. et al. Molecular mechanism of light-driven sodium pumping. Nat. Commun. 11, 2137 (2020).

51. Voss, J. M., Harder, O. F., Olshin, P. K., Drabbels, M. & Lorenz, U. J. Rapid melting and revitrification as an approach to microsecond time-resolved cryo-electron microscopy. Chem. Phys. Lett. 778, 138812 (2021).

52. Williams, H. M. et al. Overcoming Preferred Orientation in Cryo-EM With Ultrasonic Excitation During Vitrification. 2025.09.14.676144 Preprint at 10.1101/2025.09.14.676144 (2025).

53. Danev, R., Yanagisawa, H. & Kikkawa, M. Cryo-Electron Microscopy Methodology: Current Aspects and Future Directions. Trends Biochem. Sci. 44, 837–848 (2019).

