## Supplementary Information for "Microsecond Time-Resolved Cryo-EM Based on Jet Vitrification"

**This PDF file includes:**

1 | Jet vitrification experiments

2 | Capture of high-speed video frames of the jet vitrification process

3 | Estimate of the time resolution of the experiment

4 | Prediction of the arrival time of the jet

5 | Cryo-EM sample preparation

6 | Data collection and analysis – apoferritin

7 | Data collection and analysis – *Er*NaR

8 | References

### 1 | Jet vitrification experiments

Figure S1 illustrates details of the jetting assembly in a cutaway view. The activation subunit (left), which is held at 290 K, and jetting subunit (right), cooled to 95 K, are both machined from brass, which provides high thermal conductivity. They are connected to each other with two stainless steel rods, which limits the heat transfer between the subunits. The entire jetting assembly is sandwiched between two 15 mm thick half-shells machined from thermally insulating Airex T90 closed-cell foam and is mounted underneath the climate chamber of the plunge freezing device (Fig. 1c).

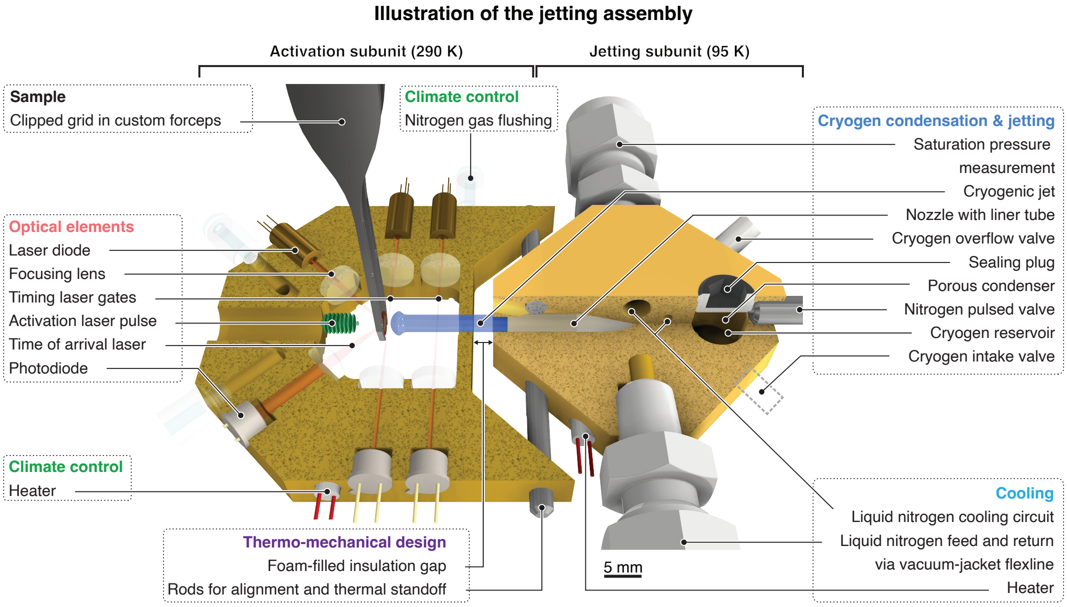

**Figure S1 | Illustration of the jetting assembly.**

**The jetting subunit.** The jetting subunit is cooled with liquid nitrogen, which is delivered through a vacuum-jacketed coaxial flexline. The flexline is pressurized to 5 bar, which raises the boiling point of nitrogen to about 94 K, above the freezing point of ethane (about 90 K at atmospheric pressure). A Lakeshore 336 controller stabilizes the temperature to 95 ± 1 K, using a cartridge heater and Pt1000 temperature sensor that are embedded in the subunit. In order to create a cryogenic jet, approximately 0.3 mL of liquid ethane (N45 purity, or alternatively, a 1:1 ethane/propane mixture with N45 and N35 purity, respectively) are condensed into a reservoir within the jetting subunit. To this end, two valves on each side of the jetting unit are opened (BMT AMV-ENM-24-01), allowing ethane to flow through a porous plug of sintered metal beads that condenses the gas into the reservoir. After 60 s, the valves are closed. Finally, we create a jet of the liquid cryogen by expelling it from the reservoir with a pulse of compressed nitrogen gas (typically 650 μs, 22 bar, N50 purity), which is applied with a Parker pulsed valve (#009-0582-900) controlled by an Iota One driver. The liquid jet exits the jetting unit through a circular channel that is lined with a smooth PTFE tube (29 mm long, 2 mm inner diameter) and travels toward the sample with a typical speed of 2.6 m/s. We do not find a significant difference between using pure ethane jets or jets with a 1:1 ethane/propane mixture.[1–4] Note that we reject experiments in which no stable jet was generated, which can for example be detected with the timing signals that are described below. For a skilled operator, at least 90 % of the experiments yield usable jets. Prior to condensing more cryogen into the reservoir and generating another jet, we purge the jetting unit with compressed nitrogen gas.

**The activation subunit.** For a time-resolved jet vitrification experiment, a clipped sample, held by custom forceps (Fig. 1c), is placed in the activation subunit, where it is illuminated with a laser pulse from the left, and vitrified with a cryogenic jet that is generated in the jetting subunit on its right. The temperature of the activation subunit is stabilized to 290 ± 1 K with a Lakeshore 336 controller, using a cartridge heater and Pt1000 temperature sensor. To prevent condensation on any of the elements, the subunit is continuously purged with a stream of dry nitrogen. On its way to the sample, the passage of the cryogenic jet is detected with two laser gates, each consisting of a laser beam that traverses the subunit and that is deflected by the passage of the jet. The output of a 650 nm laser diode (LC-LMD-650) is focused by a 10 mm lens (Thorlabs LA1116-A) to a waist of about 40 μm FWHM in the center of the activation subunit (as measured with a knife edge scan) and detected on the opposite side by a photodiode (Osram BPX 65, 1 $mm^2$ active area). The photodiode signals are amplified, and a drop is detected to provide trigger signals, which are then used by an FPGA (Digilent Cmod A7-35T) to predict the arrival time of the jet on the fly (Supplementary Information 4). A third laser beam is used to detect when the jet impinges on the sample. To this end, the output from another 650 nm laser diode is focused and reflected off the back of the sample at an angle of 45°. The arrival of the jet deflects the beam, which is similarly detected with a photodiode. The amplified signals of all photodiodes are recorded with an oscilloscope (LeCroy WaveSurfer 3054), allowing us to determine the precise arrival time of the jet after the experiment.

**Integration into a commercial plunge freezing device.** As shown in Fig. 1c, the jetting assembly is integrated into a commercial plunge freezing device (Thermo Fisher Vitrobot Mark IV). Clipped specimen supports are loaded into stainless steel precision forceps (Fig. 1c, inset) that we modified to accept clipped grids. The forceps are attached to the plunge rod of the plunge freezing device, and a sample is prepared in the climate chamber by applying the sample solution to the specimen support, followed by single-sided blotting. Once the blotting arms have returned to their initial position, which is detected by a limit switch, custom electronics take control of the stepper motor that moves the plunge rod. The sample is lowered into the jetting assembly, which is mounted underneath the climate chamber of the plunge freezing device. In order to improve the reproducibility with which the sample can be positioned in the jetting assembly, we added another linear bearing to better guide the vertical movement of the plunge rod. The stepper motor driver (TMCM-1110) is operated in microstepping mode, allowing us to position the sample in the activation subunit with a vertical resolution of about 24 µm. After the sample has been positioned in the jetting assembly, we wait for at least 200 ms in order for oscillations of the plunge rod and forceps to subside, after which the time-resolved jet vitrification experiment is performed. The sample is then immediately lowered into a liquid ethane/propane filled brass cup placed underneath the jetting assembly, from where it is transferred for imaging.

**The activation laser.** We initiate dynamics with a nanosecond laser pulse (6 ns FWHM pulse duration) with a wavelength of 532 nm. The nanosecond laser (Innolas Spitlight 600) operates at a repetition rate of 20 Hz. In the experiment, the flash lamps of the laser are therefore continuously pulsed at 20 Hz, while the Q-switch is activated only once to extract a single laser pulse and initiate dynamics. The triggers for the flash lamps and Q-switch are provided by the FPGA that controls the timing of the experiment. The Gaussian laser beam is expanded to a size of about 10 mm FWHM at the sample location, which yields a fluence variation across the viewing area of the specimen support of less than 5 %. The fluence at the sample location is adjusted with a half wave plate and polarizer.

**2 | Capture of high-speed video frames of the jet vitrification process**

The video frames of a jet impinging on a specimen support in Fig. 2a are captured with a high-speed camera operating at a frame rate of 7900 frames/s (Chronos 1.4, Kron Technologies). The camera views the sample and the jet from below the jetting assembly. We determine when the jet impinges on the sample from the video frames and calculate the approximate time stamps of the video frames indicated in Fig. 2a accordingly.

**3 | Estimate of the time resolution of the experiment**

The time resolution of our experiment is determined by the speed with which we can jet vitrify the sample to trap proteins in their transient configurations as well as the accuracy with which we can determine at which moment vitrification occurs. We estimate how fast a sample cools during jet vitrification by placing a resistance temperature detector at the sample location (Wildfire Nano-Chip, DENS Solutions DENS-P-U-H-SS-1). The sensing area of the detector consists of a 150 nm thick molybdenum spiral with a diameter of about 175 μm on a 400 nm thick silicon nitride substrate. The detector, operated in a 4-wire arrangement, is subjected to a two-point calibration (295 K and 77 K), and read out with a fast oscilloscope (LeCroy WaveSurfer 3054). Figure S2 displays a typical temperature evolution (red) during the impact of an ethane-propane jet (1:1 mixture) traveling at 2.65 m/s. As the jet impinges on the sample, the temperature drops rapidly from room temperature to below 100 K. An exponential fit (solid, black line) yields a 1/e cooling time of 15.7 μs. We simultaneously record the signal from the photodiode that detects the arrival time laser (blue, Fig. 1b), which is reflected off the back of the temperature detector. As the jet impinges on the detector, the laser beam is deflected, causing the photodiode signal to drop. We determine the timing of this event from the intersection of two linear fits to the signal (dashed, black lines). Note that the onset of cooling as determined from the exponential fit of the temperature evolution occurs 9.2 μs later. This is because the signal from the temperature detector initially drops more gradually than the exponential fit.

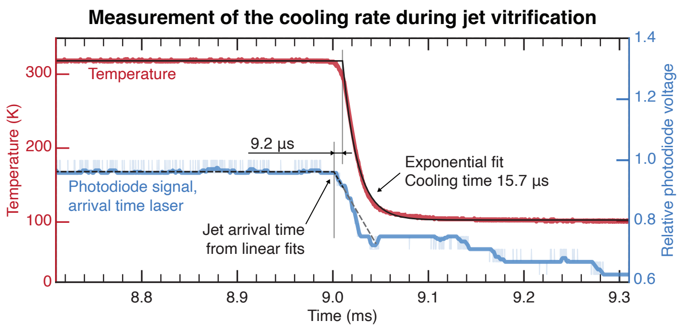

**Figure S2 | Measurement of the cooling rate during jet vitrification (from Fig. 2b).** Typical temperature evolution (red) during the impact of the jet (1:1 ethane-propane, traveling at 2.65 m/s), as measured with a resistance temperature detector (Wildfire Nano-Chip) placed at the location of the sample. An exponential fit (solid black line) yields a 1/e cooling time of 15.7 µs. The impact of the jet is detected with the arrival time laser, which is reflected off the back of temperature detector. As the jet impinges on the detector, the laser beam is deflected, which is detected with a photodiode (light blue, the dark blue curve represents the smoothed signal). We determine the timing of this event from the intersection of two linear fits to the signal shown as dashed, black lines (with the dashed, grey line to guide the eye). The time is indicated with respect to the trigger sent to the pulsed valve, which releases a pulse of compressed nitrogen that expels the liquid jet.

From a total of 18 ethane jets, we obtain and average cooling time of 19 ± 8 µs as well as an average delay between impact of the jet as obtained from the photodiode signal and the onset of cooling as determined from the exponential fit of -7 ± 9 µs. This allows us to estimate that our experiment has a time resolution of 21 µs, as obtained from the root sum square of the average cooling time and the uncertainty of the detection of the onset of cooling with the arrival time laser.

Note that the detector measures the average temperature evolution within an area of about 175 µm diameter. Different regions within this area likely cool at different times, depending on when the jet reaches the region. Within such a small region, the detector cools much faster, with a time constant on the order of 5 µs, as we can estimate from heat transfer simulations. Because of its lower heat capacity, the simulated time constant for an actual cryo-EM sample is about 5 times faster. Therefore, the measurement with the resistance temperature detector likely provides a conservative estimate for the maximum time resolution of our experiment. Also note that because of the curvature of the jet front (Fig. 2a), areas further away from the center of the grid are vitrified later. We can conservatively estimate this time delay if we calculate when the jet would arrive on different parts on the grid, based on a typical jet speed of 2.6 m/s and radius of curvature in the center of the jet of 1.8 mm. In the experiment in Fig. 3, we have collected data from within a radius of less than 400 μm from the center of the grid to obtain a transient structure of *Er*NaR at a time delay of 107 μs. While the timing spread encountered within the 175 μm diameter sensing area of the detector is contained in our measurement of the cooling time, collecting data from this larger area adds a timing uncertainty of about 14 μs.

**4 | Prediction of the arrival time of the jet**

We synchronize the activation laser pulse to the predicted arrival time of the jet. Depending on the desired time delay, we employ a different method of synchronization, which allows us to obtain an improved precision for shorter time delays. All events are timed with an FPGA (Digilent Cmod A7-35T), which provides a timing resolution of 8 ns.

**Time delays longer than 3 ms.** For the longest time delays, we simply synchronize the activation laser to the pulse of compressed nitrogen that expels the cryogenic jet. Figure S3 illustrates a typical experiment, showing the signal recorded with the photodiode that monitors the arrival time laser (Fig. 1b). As the jet impinges on the sample, the laser beam is deflected, which causes the signal to drop at 11.1 ms (as determined from the intersection of two linear fits, see above). The signal also shows a sharp positive spike at 0.7 ms, which arises from the activation laser pulse, which is scattered by the sample. Here, we have set the laser pulse to arrive 10 ms before the jet. Due to the uncertainty in the arrival time of the jet, the actual time delay is 10.4 ms. From a total of 64 such experiments, we determine that the jet impinges on the sample at an average time delay of 10.7 ms, with a standard deviation of 0.78 ms, which determines the uncertainty with which we can predict the arrival time of the jet when we synchronize the laser pulse to the trigger provided to the pulsed valve.

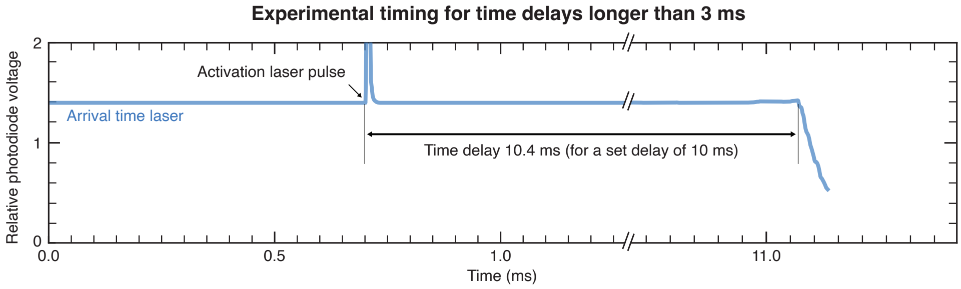

**Figure S3 | Experimental timing for time delays longer than 3 ms.** For long time delays, the activation laser is synchronized to the pulse of compressed nitrogen that expels that cryogenic jet. We detect the impact of the jet with the arrival time laser, which is reflected off the back of the sample. As the jet impinges on the sample, the laser beam is deflected, which is detected with a photodiode (blue).

**Intermediate time delays of 0.3–3 ms.** For the intermediate time delays, we synchronize the activation laser to the pulse to the passage of the cryogenic jet through the first laser gate (Fig. 1b). Figure S4 illustrates a typical experiment. As the jet passes through the gate, the laser beam is deflected, which we monitor with a photodiode (orange). An electronic circuit detects the signal drop, which is then used to time the activation laser. As above, the arrival of the jet on the sample is monitored with the arrival time laser (blue). Here, we have set the activation laser pulse to arrive 2 ms before the jet, with the actual time delay being 1.97 ms. From a total of 64 such experiments, we determine that on average, the jet impinges on the sample 3.1 ms after passing through the first laser gate, with a standard deviation of 0.39 ms, which determines the uncertainty with which we can predict the arrival time of the jet. In our analysis, we determine the time at which the jet passes through the first gate from a 30 % signal drop. The moment the jet impinges on the sample is determined from the arrival time laser signal, using the intersection of two linear fits to describe the signal drop.

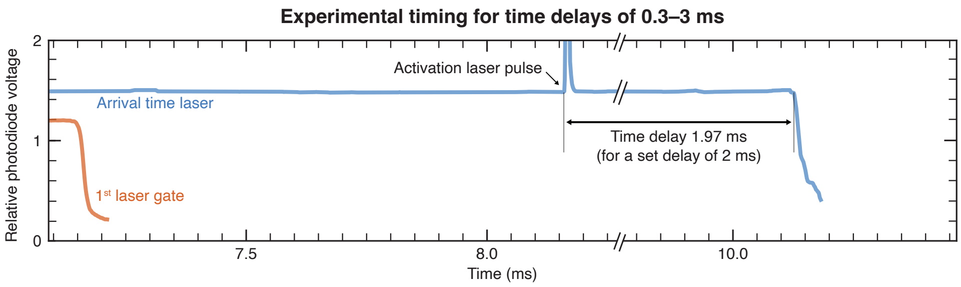

**Figure S4 | Experimental timing for time delays of 0.3–3 ms.** For intermediate time delays, the activation laser is synchronized to the passage of the cryogenic jet through the first laser gate. The jet deflects the laser beam of the gate, which we detect with a photodiode (orange). The arrival of the jet on the sample is detected with the arrival time laser, which is reflected off the back of the sample. As the jet impinges on the sample, the laser beam is deflected, which is detected with a photodiode (blue).

**Time delays shorter than 300 µs.** For the shortest time delays, we predict the arrival time of the jet on the fly by monitoring the passage of the jet through the two laser gates (Fig. 1b). This then allows us to synchronize the activation laser accordingly. Figure S5 illustrates a typical experiment. The signals from the photodiodes that monitor the passage of the jet through the laser gates are shown in orange and magenta. An electronic circuit detects the signal drop. This timing information is then used by the FPGA to predict the arrival time of the jet on the fly. Figure S5 also shows an additional digital signal generated by the FPGA that marks the predicted arrival time (green). As above, the actual arrival of the jet is monitored with the arrival time laser (blue). Here, we have set the activation laser pulse to arrive 100 µs before the jet, with the actual time delay being 119 µs.

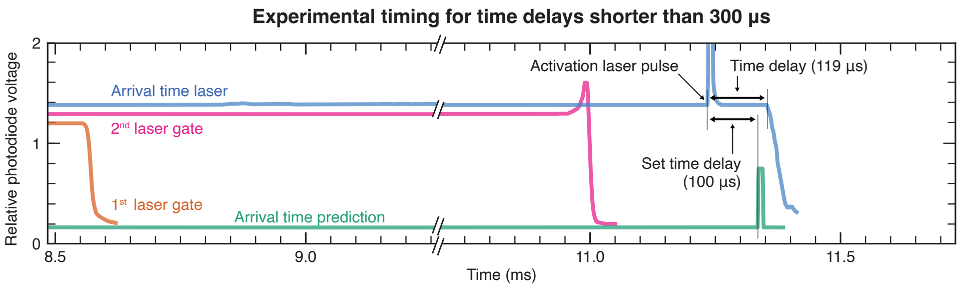

**Figure S5 | Experimental timing for time delays shorter than 300 µs.** For the shortest time delays, we predict the arrival time of the cryogenic jet on the sample on the fly from the passage of the jet through both laser gates and time the activation laser pulse accordingly. The passage of the jet through the gates deflects the laser beam, which we detect with photodiodes (orange and magenta). The arrival time of the jet on the sample is then calculated on the fly with an FPGA, which is used to time the laser pulse. Here, we also show an additional digital signal generated by the FPGA that marks the predicted arrival time (green). The actual arrival of the jet is detected with the arrival time laser, which is reflected off the back of the sample. As the jet impinges on the sample, the laser beam is deflected, which is detected with a photodiode (blue).

The FPGA predicts the arrival time of the jet with the following method. From the timing information provided by the two laser gates, which are placed at a distance of 7 mm, we can calculate the jet speed. With an effective distance of the sample from the second laser gate of about 0.92 mm, we can then predict the arrival time of the jet. Before conducting time-resolved experiments, we first record the timing information for several test experiments, from which we then derive the effective distance, which we take to be the average of the distance calculated in each experiment. Note that this effective distance, for which our calculation provides the most accurate prediction of the arrival time, differs slightly from the actual distance. One reason for this discrepancy is that our calculation assumes that the jet velocity is constant, whereas the jet slightly accelerates on its way to the sample.

From a total of 64 experiments, we obtain an average jet speed between the gates of 2.6 ± 0.4 m/s and an effective distance of the sample from the second gate of 0.92 ± 0.11 mm. In our experiment, the average time delay between the predicted arrival time of the jet, as determined from the FPGA output, and the actual arrival time is -22 µs, with a standard deviation of 38 µs (16 experiments). The standard deviation provides a measure of the uncertainty of the prediction of the jet arrival time. In our analysis, we determine the time at which the jet passes through the gate from a 30 % signal drop. The predicted arrival time is determined from the digital timing signal provided by the FPGA (green trace in Fig. S5), and the moment the jet impinges on the sample is determined from the arrival time laser signal. For both signals, we use the intersection of two linear fits to obtain the precise timing of the event.

**5 | Cryo-EM sample preparation**

Mouse heavy chain apoferritin (5 mg/mL, buffer 10 mM HEPES pH 7.5, 150 mM NaCl) was provided by the Protein Production and Structure Core Facility (PTPSP) at EPFL, Lausanne, Switzerland. Holey gold grids (R2/2, 200 gold mesh, Quantifoil) were clipped (with the top side facing towards the clip ring and the bar side towards the c-clip, Sub-Angstrom) and rendered hydrophilic in a glow discharge (air, 90 s duration, 15 mA current, EasiGlow, Ted Pella). The cryo-EM sample was prepared by applying 3 µL sample solution to the grid, followed by single-sided blotting (temperature 4 °C, 95 % relative humidity, blotting force 10, blotting time 9 s, delay before blotting 0.5 s, drain time 0.5 s). The grid was then lowered into the jetting assembly and vitrified with a jet of liquid ethane. Finally, the sample was plunged into a 1:1 liquid ethane/propane mixture and transferred to liquid nitrogen for storage until data collection.

Cryo-EM samples of *Er*NaR were prepared at a concentration of 4.5 mg/mL in 10 mM Tris-HCl, pH 8.0, 0.04 % (v/v) DDM, with a NaCl concentration of 200 mM for the dark state and 107 µs samples, and 500 mM for the 13.3 ms sample. The holey amorphous carbon sample supports (R1.2/1.3, 200 gold mesh, Quantifoil) were coated with a 40 nm thick layer of silver (Quorum Q300T) on the bottom side that is illuminated by the laser pulse, so as to reduce the amount of light absorbed by the grid and limit the associated temperature jump of the sample. The grids were clipped (with the top side facing towards the clip ring and the bar side towards the c-clip, Sub-Angstrom) and rendered hydrophilic in a glow discharge (air, 90 s duration, 15 mA current, EasiGlow, Ted Pella). The cryo-EM samples were prepared by applying 3.5 µL sample solution to the grid, followed by single-sided blotting (temperature 4 °C, 95% relative humidity, blotting force 10, blotting time 6-7 s, delay before blotting 0.5 s, drain time 0.5 s). The grids were then lowered into the jetting assembly where they were laser-activated (except for the dark state) and jet vitrified with a 1:1 liquid ethane/propane mixture. Finally, the grids were plunged into a 1:1 liquid ethane/propane mixture and transferred to liquid nitrogen for storage until data collection.

For the time-resolved experiments, the samples are excited with a nanosecond laser pulse prior to vitrification (532 nm wavelength, 6 ns FWHM pulse duration, 6.4 mJ/cm$^2$ fluence). We estimate from a multilayer optical calculation[5] that the thin film of the sample absorbs about 10 % of the laser fluence, leading to an temperature jump of about 20 K for a typical sample thickness. The temperature jump only lasts for the duration of a few microseconds as the heat is quickly dissipated towards the massive grid bars, which act as heat sink.[6] With the absorption cross section of a single rhodopsin monomer at 532 nm of $1.8 \cdot 10^{-16}$ $cm^2$ (as measured with a NanoDrop One Spectrophotometer), we saturate the transition about 3-fold.

**6 | Data collection and analysis — apoferritin**

High-resolution micrographs of a cryo-EM sample of apoferritin were collected in Dubochet Center for Imaging in Lausanne (DCI-Lausanne), using a Thermo Fisher Titan Krios G4i transmission electron microscope equipped with SelectrisX energy filter and Falcon IV direct electron detector. Thermo Fisher EPU software was used for data acquisition. Single-particle reconstructions were performed using CryoSPARC v4.6.0[7], with the corresponding workflow illustrated in Fig. S6. The data acquisition parameters and processing statistics are summarized in SI Table 1.

Micrographs were subjected to patch motion correction and patch CTF estimation. Micrographs with an estimated resolution of worse than 5 Å were rejected. Particles were picked using a blob picker with a diameter ranging from 100 to 130 Å. The particles were extracted with a box size of 720 px and Fourier-cropped to 360 px.

Particles were first subjected to 2D classification followed by *ab initio* reconstruction (one class, $C_1$ symmetry) and homogeneous refinement (O symmetry). This refined volume and five decoy volumes were then used in a single round of heterogeneous refinement (six classes, $C_1$ symmetry). Particles from the best class were subjected to local and global CTF refinement before being re-extracted with a box size of 686 px. These particles were then homogeneously refined (O symmetry) and subjected to reference-based motion correction. A final homogeneous refinement (O symmetry) was then performed producing the volume shown in Fig. S6.

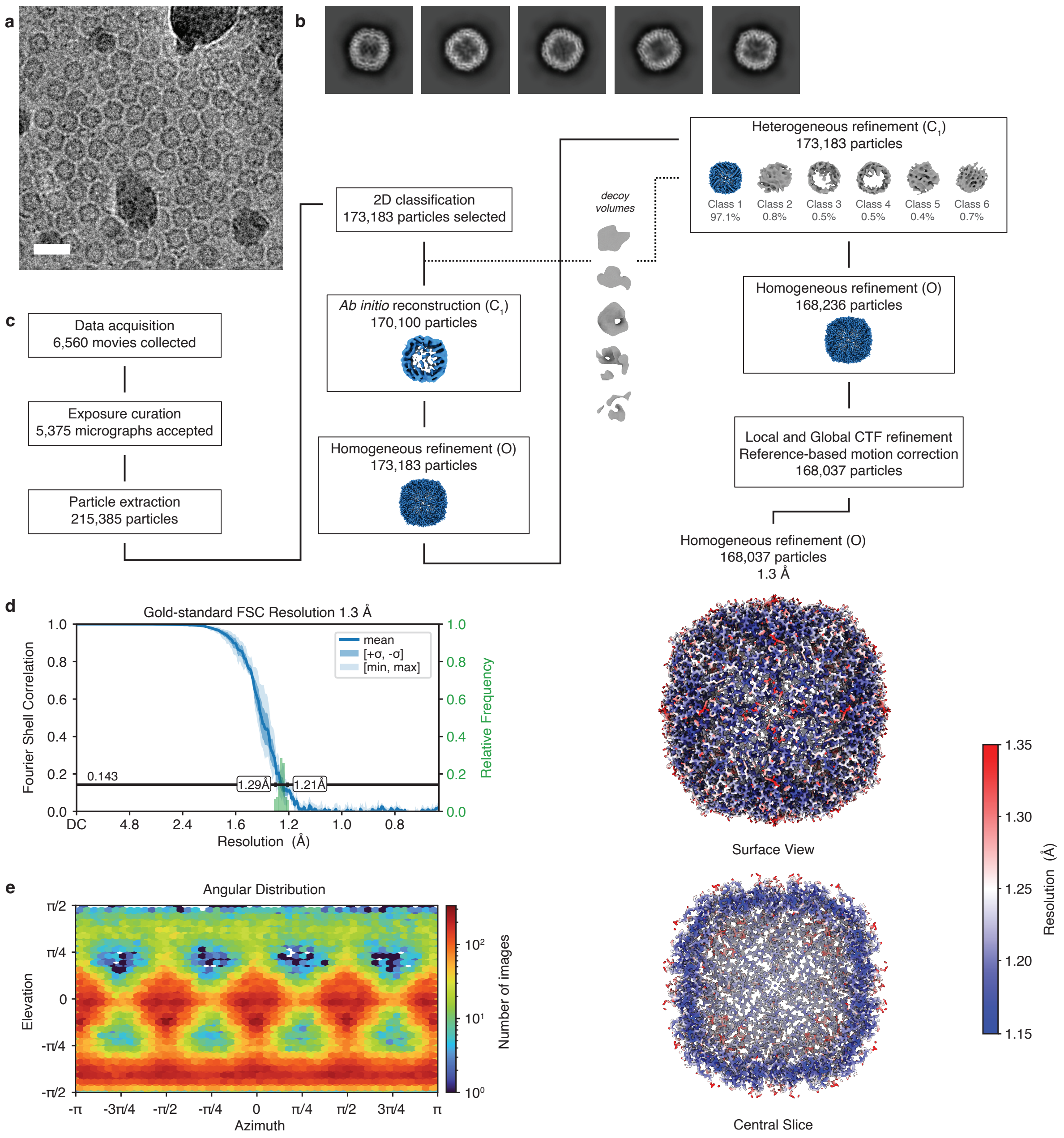

**Figure S6 | Processing workflow for the Apoferritin sample. a** Representative micrograph. Scale bar, 200 Å. **b** Selected 2D class averages. **c** Data processing workflow in CryoSPARC. Symmetry applied is indicated in parentheses. The volumes are displayed at a level of 5σ above the mean. The final map is displayed at a level 10σ above the mean, with the local resolution estimation at the FSC cutoff of 0.5 indicated in color. **d** Conical FSC, with the 0.143 threshold indicated by a black line. The mean FSC value is shown as a solid blue line. Dark blue shading indicates one standard deviation of the FSC curve, while the light blue shading represents the minimum and maximum FSC values. A histogram of the resolution values obtained from the conical FSC is shown in green. **e** Angular distribution of the particles from the final reconstruction.

**Table S1 | Cryo-EM data collection, refinement and validation statistics for apoferritin**

| | Apoferritin (EMDB-55761) |
|---|---|
| **Data collection and processing** | |
| Microscope | Titan Krios G4i |
| Camera | Falcon 4i |
| Energy filter (slit width) | SelectrisX (10 eV) |
| Magnification | 350’000 |
| Voltage (kV) | 300 |
| Electron exposure ($e^{-}/Å^2$) | 40 |
| Defocus range (µm) | -0.4 to -1.2 |
| Pixel size (Å) | 0.352 |
| Micrographs collected (no.) | 6’560 |
| Micrographs accepted (no.) | 5’375 |
| Initial particle images (no.) | 215’385 |
| Final particle images (no.) | 168’037 |
| Symmetry imposed | O |
| Map resolution (Å) | 1.3 |
| FSC threshold | 0.143 |
| Map resolution range (Å) | 1.29 – 1.21 |
| Map sharpening *B* factor ($Å^2$) | 21.5 |

**7 | Data collection and analysis — *Er*NaR**

High-resolution micrographs of *Er*NaR cryo-EM samples before activation (dark state) and 107 µs and 13.3 ms post-activation were collected in Dubochet Center for Imaging in Lausanne (DCI-Lausanne), using a Thermo Fisher Titan Krios G4i transmission electron microscope equipped with SelectrisX energy filter and Falcon IV direct electron detector. Thermo Fisher EPU software was used for data acquisition. Single-particle reconstructions were performed using a combination of CryoSPARC v4.6.0 and CryoSPARC v4.7.0[7], with the corresponding workflow illustrated Figs. S7–S9. The data acquisition parameters and processing statistics are summarized in SI Table 2.

Micrographs were subjected to patch motion correction and patch CTF estimation. Micrographs with an estimated resolution of worse than 8 Å (dark state and 13.3 ms post-activation) or 10 Å (107 µs post-activation) were rejected. Particles were picked using a combination blob picking (diameter ranging from 80 to 140 Å) and template picking. The particles were extracted with a box size of 384 px and Fourier-cropped to 192 px.

For the dark state dataset, particles were first subjected to two rounds of 2D classification, followed by *ab initio* reconstruction (three classes, $C_1$ symmetry). Particles from the best two *ab initio* classes were subjected to two rounds of heterogeneous refinement (two classes, $C_5$ symmetry) and a further round of 2D classification. This produced a particle stack containing 128,675 particles. These particles were then subjected to further *ab initio* reconstruction (three classes, $C_1$ symmetry). Particles from the best *ab initio* class were then non-uniformly refined ($C_5$ symmetry). This refined volume and three decoys were then used in a single round of heterogeneous refinement (four classes, $C_5$ symmetry). Particles from the best class were subjected to non-uniform refinement, local and global CTF refinement, reference-based motion correction, and re-extracted with a box size of 384 px. A final homogeneous refinement ($C_5$ symmetry) was then performed producing the volume shown in Fig. S7.

For the 107 µs dataset, particles were first subjected to two rounds of 2D classification, followed by *ab initio* reconstruction (three classes, $C_1$ symmetry). The best volume from the *ab initio* reconstruction job was then used with two decoy volumes in two rounds of heterogeneous refinement (three classes, $C_1$ symmetry). The best volume from the latest heterogeneous refinement job was then used in two subsequent rounds of heterogeneous refinement with two decoys (three classes, $C_1$ symmetry). Particles from the best volume were then subjected to local and global CTF refinement, reference-based motion correction, and non-uniform refinement ($C_5$ symmetry) producing a consensus refinement of the *Er*NaR pentamer. The particles were then $C_5$ symmetry expanded and subjected to 3D classification (ten classes, $C_1$ symmetry, focus mask covering one monomer). Particles belonging to classes revealing the photoisomerization of the retinal Schiff base were retained and subjected to another round of 3D classification (two classes, $C_1$ symmetry, focus mask covering one monomer). A final local refinement ($C_1$ symmetry) was then performed producing the volume shown in Fig. S8.

For the 13.3 ms dataset, particles were first subjected to two rounds of 2D classification, followed by *ab initio* reconstruction (two classes, $C_1$ symmetry), and a further round of 2D classification. Particles from the best 2D classes were then subjected to non-uniform refinement ($C_1$ symmetry) using the best *ab initio* volume. This volume was then used with one decoy in two rounds of heterogeneous refinement (two classes, $C_1$ symmetry). Particles from the best class were subjected to reference-based motion correction, 2D classification, local and global CTF correction, and then non-uniform refinement ($C_5$ symmetry). The particles were then $C_5$ symmetry expanded and subjected to 3D classification (ten classes, $C_1$ symmetry, focus mask covering one monomer). Particles belonging to classes revealing conformational dynamics of interest were retained and subjected to another round of 3D classification (three classes, $C_1$ symmetry, focus mask covering one monomer). A final local refinement ($C_1$ symmetry) was then performed producing the volume shown in Fig. S9.

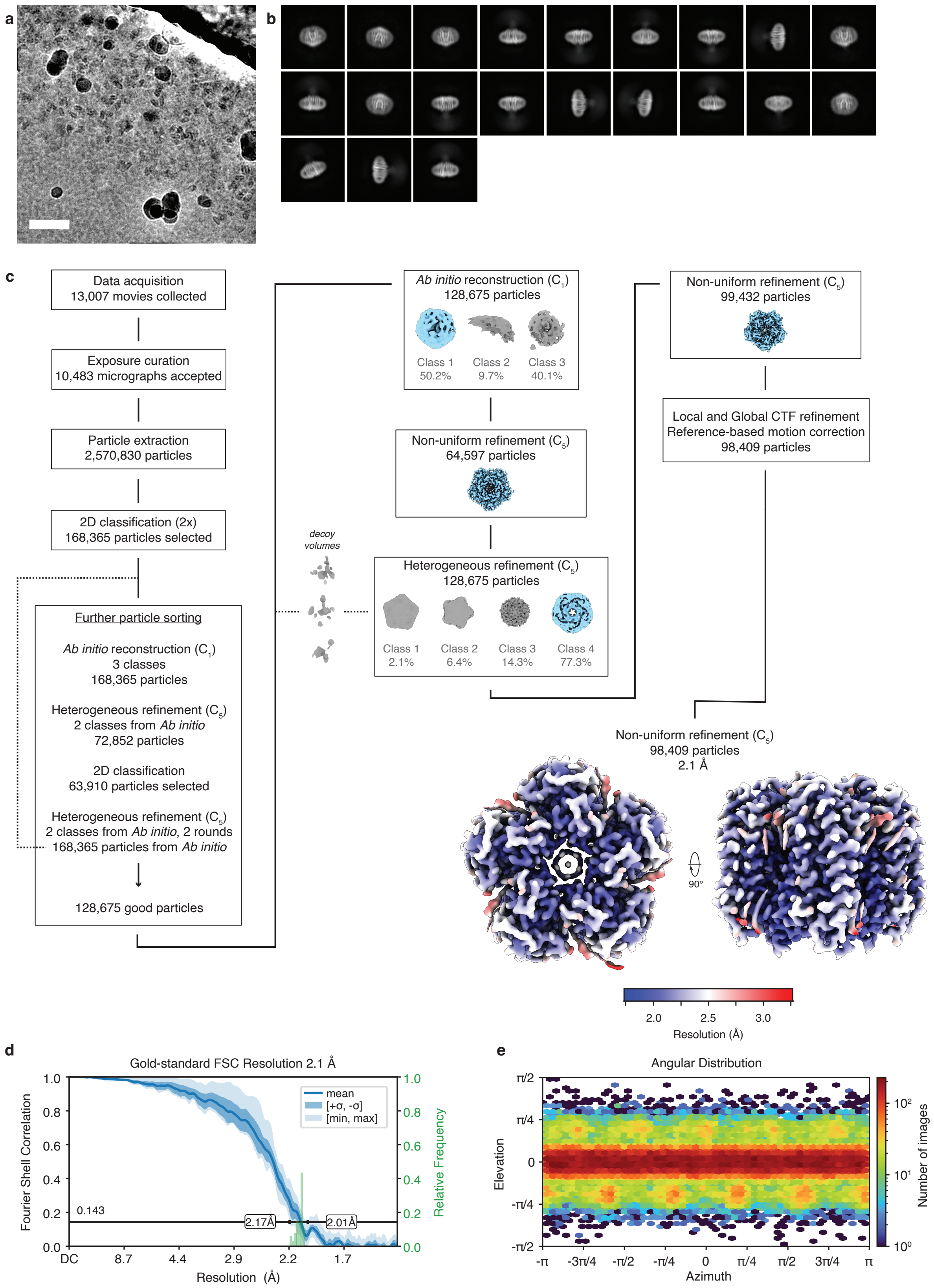

**Figure S7 | Processing workflow for the *Er*NaR dark state. a** Representative micrograph. Scale bar, 500 Å. **b** Selected 2D class averages. **c** Data processing workflow in CryoSPARC. Symmetry applied is indicated in parentheses. The volumes are displayed at a level of 10σ above the mean. The final map is displayed at a level 15$\sigma$ above the mean, with the local resolution estimation at the FSC cutoff of 0.5 indicated in color. **d** Conical FSC, with the 0.143 threshold indicated by a black line. The mean FSC value is shown as a solid blue line. Dark blue shading indicates one standard deviation of the FSC curve, while the light blue shading represents the minimum and maximum FSC values. A histogram of the resolution values obtained from the conical FSC is shown in green. **e** Angular distribution of the particles from the final reconstruction.

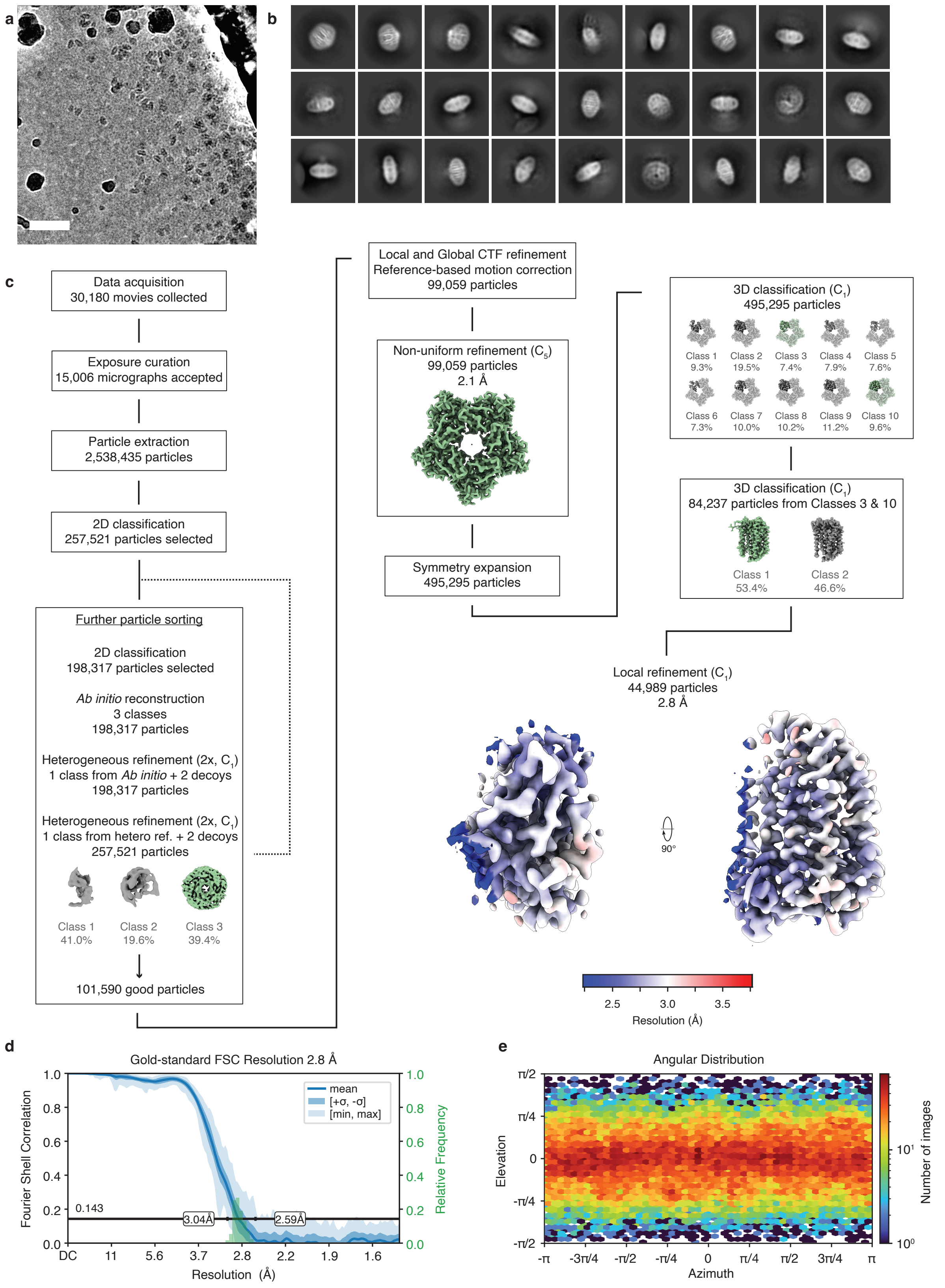

**Figure S8 | Processing workflow for the *Er*NaR photoactivated state after 107 µs.** **a** Representative micrograph. Scale bar, 500 Å. **b** Selected 2D class averages. **c** Data processing workflow in CryoSPARC. Symmetry applied is indicated in parentheses. The volumes are displayed at a level of 10σ above the mean. In 3D classification, focus mask was applied on the highlighted region. The final map is displayed at a level 15σ above the mean, with the local resolution estimation at the FSC cutoff of 0.5 indicated in color. **d** Conical FSC, with the 0.143 threshold indicated by a black line. The mean FSC value is shown as a solid blue line. Dark blue shading indicates one standard deviation of the FSC curve, while the light blue shading represents the minimum and maximum FSC values. A histogram of the resolution values obtained from the conical FSC is shown in green. **e** Angular distribution of the particles from the final reconstruction.

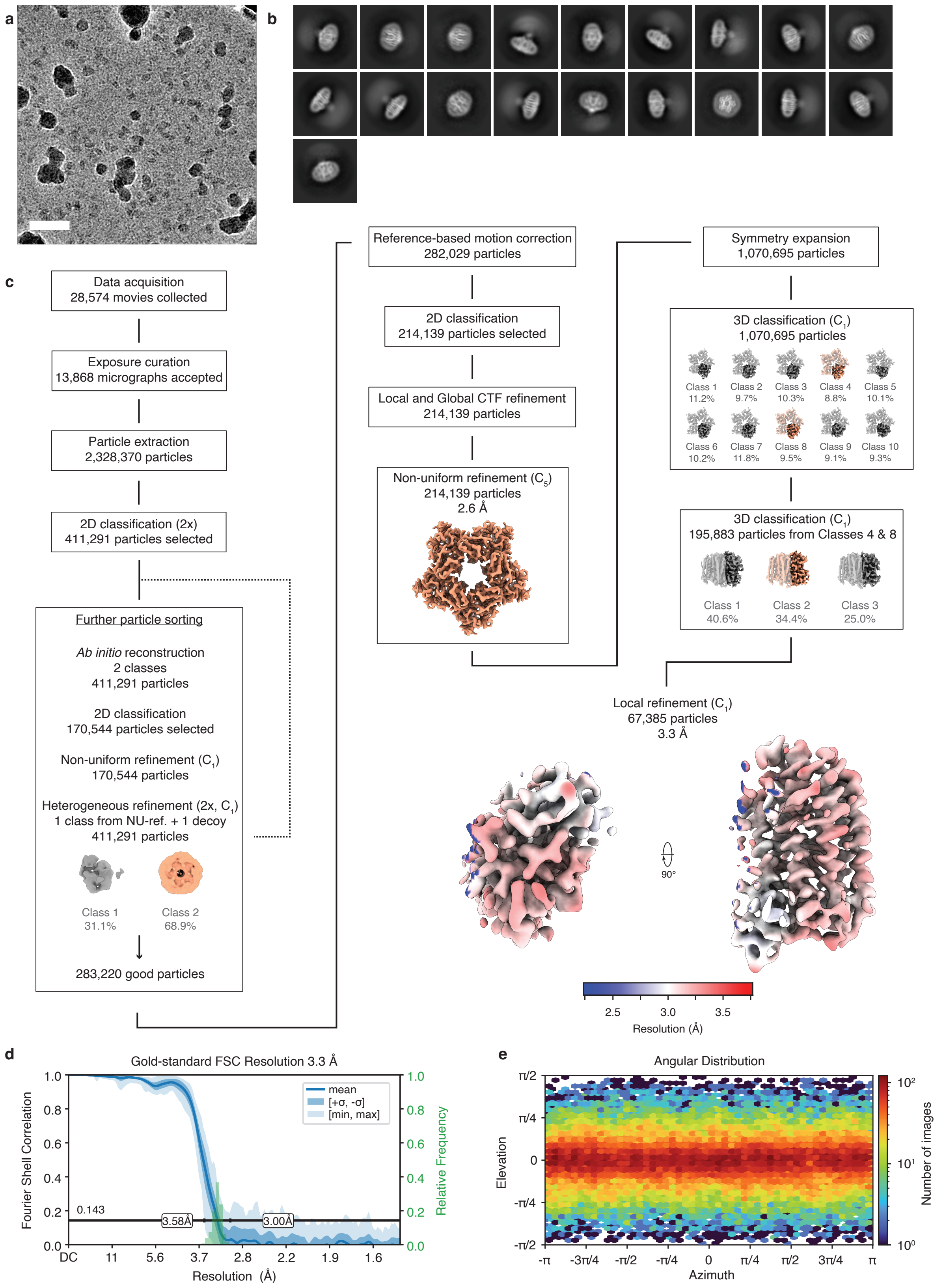

**Figure S9 | Processing workflow for the *Er*NaR photoactivated state after 13.3 ms.** **a** Representative micrograph. Scale bar, 500 Å. **b** Selected 2D class averages. **c** Data processing workflow in CryoSPARC. Symmetry applied is indicated in parentheses. The volumes are displayed at a level of 10$\sigma$ above the mean. In 3D classification, focus mask was applied on the highlighted region. The final map is displayed at a level 15$\sigma$ above the mean, with the local resolution estimation at the FSC cutoff of 0.5 indicated in color. **d** Conical FSC, with the 0.143 threshold indicated by a black line. The mean FSC value is shown as a solid blue line. Dark blue shading indicates one standard deviation of the FSC curve, while the light blue shading represents the minimum and maximum FSC values. A histogram of the resolution values obtained from the conical FSC is shown in green. **e** Angular distribution of the particles from the final reconstruction.

**Table S2 | Cryo-EM data collection, refinement and validation statistics for *Er*NaR**

| | *Er*NaR, dark state (EMDB-55765) (PDB-9TBD) | *Er*NaR, 107 μs monomer (EMDB-55766) (PDB-9TBE) | *Er*NaR, 13.3 ms monomer (EMDB-55769) (PDB-9TBF) |
|---|---|---|---|
| **Data collection and processing** | | | |
| Microscope | Titan Krios G3i | Titan Krios G3i | Titan Krios G3i |
| Camera | Falcon 4i | Falcon 4i | Falcon 4i |
| Energy Filter (slit width) | SelectrisX (10 eV) | SelectrisX (10 eV) | SelectrisX (10 eV) |
| Magnification | 165’000 | 165’000 | 165’000 |
| Voltage (kV) | 300 | 300 | 300 |
| Electron exposure ($e^{-}$/Å$^2$) | 40 | 40 | 40 |
| Defocus range (μm) | -0.8 to -2.4 | -0.8 to -2.4 | -0.8 to -2.4 |
| Pixel size (Å) | 0.726 | 0.726 | 0.726 |
| Micrographs collected (no.) | 13’007 | 30’180 | 28’574 |
| Micrographs accepted (no.) | 10’483 | 15’006 | 13’868 |
| Initial particle images (no.) | 2’570’830 | 2’538’435 | 2’328’370 |
| Final particle images (no.) | 98’409 | 44’989 | 67’385 |
| Symmetry | C5 | C1 final (C5 symmetry expanded) | C1 final (C5 symmetry expanded) |
| Map resolution (Å) | 2.1 | 2.8 | 3.3 |
| FSC threshold | 0.143 | 0.143 | 0.143 |
| Map resolution range (Å) | 2.17-2.01 | 3.04-2.59 | 3.58-3.00 |
| Map sharpening *B* factor (Å$^2$) | 49.5 | 93.7 | 127.2 |
| **Refinement** | | | |
| Initial model used (PDB code) | 8QQZ | 8QQZ | 8QQZ |
| Model composition | | | |
| Non-hydrogen atoms | 12’168 | 2’380 | 2’409 |
| Protein residues | 1’360 | 272 | 274 |
| Ligands | 56 | 13 | 13 |
| *B* factors (Å$^2$) | | | |
| Protein | 18.49 | 55.94 | 44.6 |
| Ligand | 32.07 | 69.66 | 43.56 |
| R.m.s. deviations | | | |
| Bond lengths (Å) | 0.004 | 0.004 | 0.004 |
| Bond angles (°) | 0.766 | 0.915 | 0.793 |
| Validation | | | |
| MolProbity score | 1.59 | 1.75 | 1.95 |
| Clashscore | 6.24 | 4.77 | 5.72 |
| Poor rotamers (%) | 2.09 | 3.36 | 2.71 |
| Ramachandran plot | | | |
| Favored (%) | 99.10 | 97.38 | 95.54 |
| Allowed (%) | 0.90 | 2.62 | 4.46 |
| Disallowed (%) | 0.00 | 0.00 | 0.00 |

**8 | References**

1. Ravelli, R. B. G. *et al.* Cryo-EM structures from sub-nl volumes using pin-printing and jet vitrification. *Nat. Commun.* **11**, 2563 (2020).
2. Müller, T., Moser, S., Vogt, M., Daugherty, C. & Parthasarathy, M. V. Optimization and Application of Jet-Freezing. *Scanning Microsc.* **7**, (1993).
3. Bald, W. B. The relative merits of various cooling methods. *J. Microsc.* **140**, 17–40 (1985).
4. Ryan, K. P., Purse, D. H., Robinson, S. G. & Wood, J. W. The relative efficiency of cryogens used for plunge-cooling biological specimens. *J. Microsc.* **145**, 89–96 (1987).
5. Byrnes, S. J. Multilayer optical calculations. Preprint at https://doi.org/10.48550/arXiv.1603.02720 (2020).
6. Voss, J. M., Harder, O. F., Olshin, P. K., Drabbels, M. & Lorenz, U. J. Rapid melting and revitrification as an approach to microsecond time-resolved cryo-electron microscopy. *Chem. Phys. Lett.* **778**, 138812 (2021).
7. Punjani, A., Rubinstein, J. L., Fleet, D. J. & Brubaker, M. A. cryoSPARC: algorithms for rapid unsupervised cryo-EM structure determination. *Nat. Methods* **14**, 290–296 (2017).
